# A single genetic locus lengthens deer mouse burrows via motor pattern evolution

**DOI:** 10.1101/2023.07.03.547545

**Authors:** Olivia S. Harringmeyer, Caroline K. Hu, Hillery C. Metz, Eris L. Mihelic, Charlie Rosher, Juan I. Sanguinetti-Scheck, Hopi E. Hoekstra

**Affiliations:** Department of Molecular & Cellular Biology, Department of Organismic & Evolutionary Biology, Museum of Comparative Zoology, Center for Brain Science, Howard Hughes Medical Institute, Harvard University, Cambridge, MA, USA

## Abstract

The question of how evolution builds complex behaviors has long fascinated biologists. To address this question from a genetic perspective, we capitalize on variation in innate burrowing behavior between two sister species of *Peromyscus* mice: *P. maniculatus* that construct short, simple burrows and *P. polionotus* that uniquely construct long, elaborate burrows. We identify three regions of the genome associated with differences in burrow length and then narrow in on one large-effect 12-Mb locus on chromosome 4. By introgressing the *P. polionotus* allele into a *P. maniculatus* background, we demonstrate this locus, on its own, increases burrow length by 20%. Next, by recording mice digging in a transparent tube, we find this locus has specific effects on burrowing behavior. This locus does not affect time spent digging or latency to dig, but rather affects usage of only two of the primary digging behaviors that differ between the focal species: forelimb digging, which loosens substrate, and hindlimb kicking, which powerfully ejects substrate. This locus has an especially large effect on hindkicking, explaining 56% and 22% of interspecific differences in latency and proportion of hindkicks, respectively. Together, these data provide genetic support for the hierarchical organization of complex behaviors, offering evolution the opportunity to tinker with specific behavioral components.

---

How complex behaviors evolve is a question that has long fascinated biologists. Home- building, courtship, and parental care are examples of largely innate behaviors that vary in their complexity, even between closely related taxa ^e.g., 1–3^. A long-standing hypothesis is that complex behaviors, such as these, are comprised of hierarchically organized modules, where internal states (e.g., fear or hunger) can drive the expression of numerous independent downstream motor patterns related to that state^4–9^. This framework also raises the possibility that individual genetic loci may influence specific components of complex behaviors, while leaving others unchanged. Thus, the effects of natural genetic variation may reflect the underlying organization of behavior.

To test this hypothesis, we capitalize on a behavior that has elaborated within a lineage of deer mice (genus *Peromyscus*)^10^. In particular, two closely related species show striking differences in their burrow architecture: *Peromyscus maniculatus* build short, simple burrows (mean length = 8.58±2.15 cm), whereas its sister species *P. polionotus* construct long, complex burrows (mean length = 40.0±16.5 cm), even in a controlled laboratory environment (Fig. 1a-c). This difference in species-specific burrow architecture has a strong genetic component^11–13^.

**Fig. 1.**
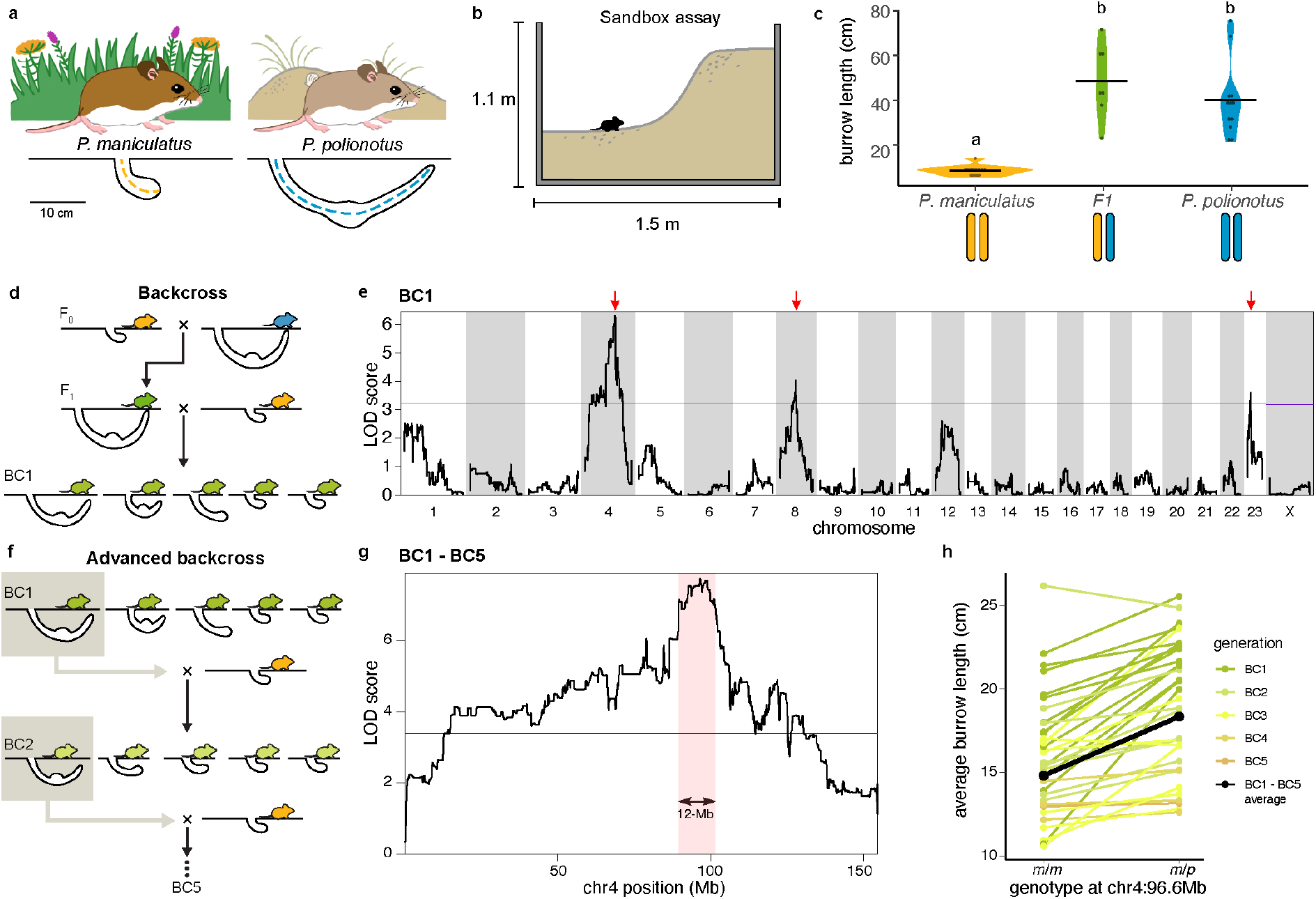
The genetic basis of innate differences in burrowing behavior. (**a**) Characteristic differences in habitat and burrow construction for two sister species of deer mice, *P. maniculatus* and *P. polionotus*. (**b**) Cross-section of large sand enclosures used to measure burrowing in the laboratory. (**c**) Burrow lengths constructed in the sand enclosures for *P. maniculatus* (yellow, *n* = 13), *P. polionotus* (blue, *n* = 12) and F1 hybrids (green, *n* = 7) with black lines denoting mean burrow length by group. Letters indicate statistically different groups (K-W test with Dunn’s test, *P* < 0.0001). (**d**) Cross design used to produce BC1 hybrids to characterize the genetic architecture of burrow length. (**e**) QTL map of log(burrow length) across BC1 hybrids (*n* = 271) using Haley-Knott regression. Significance threshold (purple line) calculated for autosomes and separately for the X-chromosome. Red arrows point to 3 genomic regions significantly associated with variation in burrow length. (**f**) Advanced backcross design with phenotypic selection to refine the QTL-chr4 region. Repeated backcrossing to *P. maniculatus* (yellow) for 5 generations (BC5). (**g**) QTL map of log(burrow length) for chr4 across BC1 – BC5 mice (*n* = 892). QTL mapping was performed using a linear mixed model with pedigree as a random effect. Significance threshold (purple line) and Bayes 95% credible interval (shaded red, 12 Mb) are given. (**h**) Effects of QTL-chr4 on burrow length for each BC family. Mice are grouped by genotype based on peak LOD-score marker at chr4:96.6-Mb (*m*/*m* = homozygous *P. maniculatus*; *m*/*p* = heterozygous *P. maniculatus*/*P. polionotus*) with points showing average burrow length of siblings from the same family, colored by BC generation. Average burrow length across all backcross mice shown in black. Statistically significant difference in burrow lengths between genotypes (linear mixed model with family as random effect, *P* < 0.001). Sample sizes: BC1: *m/m* (*n* = 141), *m/p* (*n* = 130); BC2: *m/m* (*n* = 88), *m/p* (*n* = 64); BC3: *m/m* (*n* = 220), *m/p* (*n* = 108); BC4: *m/m* (*n* = 111), *m/p* (*n* = 18); BC5: *m/m* (*n* = 5), *m/p* (*n* = 4).

While most *Peromyscus* species build short, *maniculatus*-like burrows, the length and shape of *P. polionotus* burrows are unique, representing the derived state, and thus can serve as a model for the evolution of behavioral complexity^14^.

## Genetic mapping of burrowing behavior locus

To characterize genetic loci contributing to variation in burrowing behavior, we performed quantitative trait locus (QTL) mapping of naturalistic burrowing using a backcross strategy to maximize power under dominant inheritance (Fig. 1c). We initially revisited a previous mapping population that was created by backcrossing F1 hybrids to *P. maniculatus* to produce first-generation backcross (BC1) hybrids (Fig. 1d)^12^. Burrowing was quantified for each mouse in a large sand-filled enclosure (Fig. 1b), with polyurethane casts used to measure each burrow. We re-sequenced these BC1 mice and used a chromosome-level genome assembly for *P. maniculatus* to genotype these mice at approximately 100,000 genome-wide markers (see Methods). Using QTL mapping, we identified three regions of the genome showing significant associations with variation in total burrow length (Fig. 1e). Two of these regions overlapped with those previously reported for length of only the entrance tunnel^12^: QTL on chromosomes 4 and 23, with *P. polionotus* ancestry increasing total burrow length by an average of 4.4 and 3.4 cm respectively (Extended Data Fig. 1). The chromosome 4 QTL (hereafter referred to as “QTL- chr4”), which had a 95% Bayes credible interval spanning 30.3 Mb (ranging from chr4: 77.7 – 108.0 Mb), showed the strongest effect on burrow length, explaining 10.2% of the variance among BC1 hybrids (Extended Data Fig. 1).

To refine this QTL-chr4 interval to a narrower region of the chromosome, we performed an advanced backcross. Specifically, *P. polionotus* was backcrossed to *P. maniculatus* for six generations (to create five BC generations), using phenotypic selection for long burrows at each generation (Fig. 1f, Extended Data Fig. 2; see Methods). QTL mapping of the advanced BC population (see Methods) refined QTL-chr4 by >60% to a 95% Bayes credible interval spanning 12.0 Mb (chr4: 89.6 – 101.6 Mb) (Fig. 1g). *P. polionotus* ancestry at this 12Mb locus increased burrow lengths by an average of 3.6 cm across all backcross mice (Fig. 1h).

**Fig. 2.**
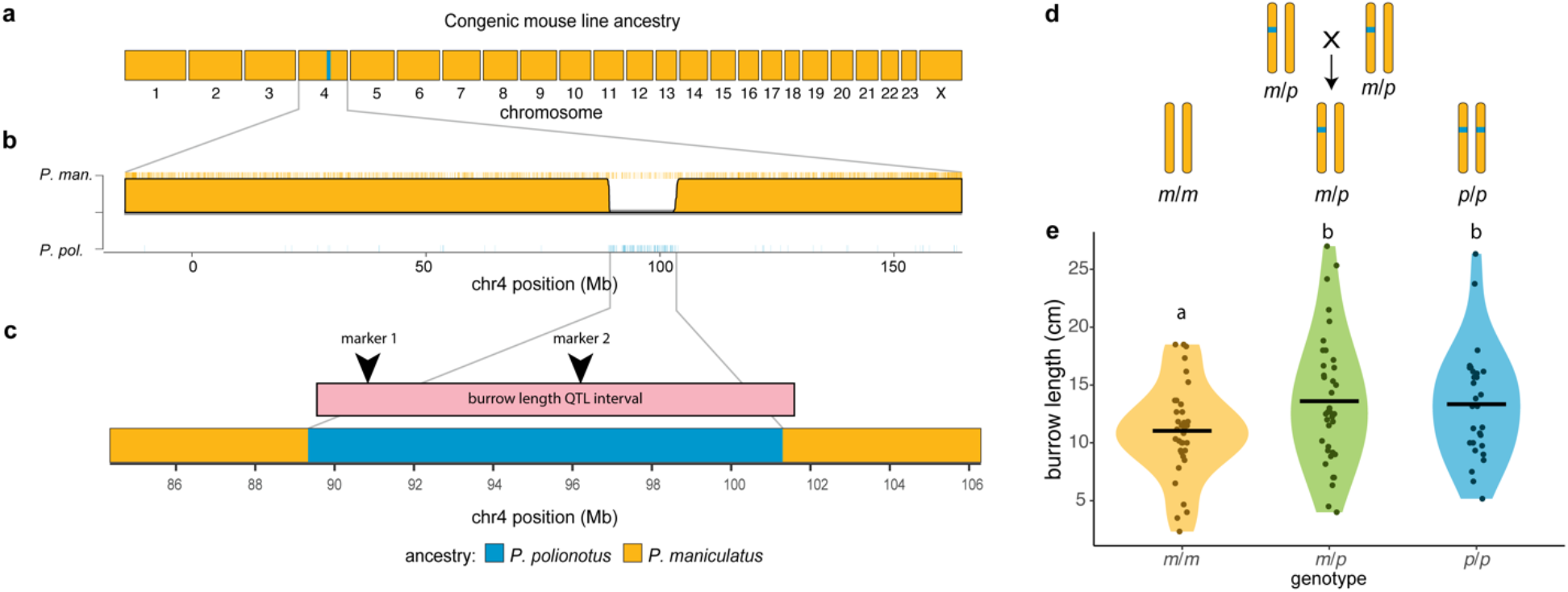
Isolating the chr4 locus in a congenic mouse line. (**a**) Genome-wide ancestry in the congenic line is shown, with *P. maniculatus* ancestry (yellow) except at chromosome 4 (*P. polionotus*, blue). (**b**) Chromosome 4 ancestry probabilities for the congenic line founder. Tick marks represent ancestry informative SNPs. (**c**) The 95% Bayes credible interval from fine- mapping (pink, above) in comparison to estimated ancestry breakpoints on chromosome 4 in the congenic line (below). Two markers used for genotyping are shown (arrows). (**d**) Breeding design for congenic line: heterozygous mice were interbred to create siblings with the three QTL-chr4 genotypes (*m*/*m* = homozygous *P. maniculatus*; *m*/*p* = heterozygous; *p*/*p* = homozygous *P. polionotus*). (**e**) Effect of QTL-chr4 on burrow length for each congenic genotype with black lines denoting mean burrow length by genotype. Sample sizes: *m/m* (*n* = 35), *m/p* (*n* = 38), *p/p* (*n* = 29). Letters indicate statistically different groups (two-sided t-tests, *P* < 0.05).

## Generating a congenic line for QTL-chr4

To isolate the implicated region on a common genomic background, we next introgressed this narrow *P. polionotus* QTL-chr4 allele into a *P. maniculatus* genome (i.e., generated a congenic line) using successive backcrossing paired with molecular genotyping. This strategy provided a uniform genetic background on which to test the specific effects of the introgressed locus, eliminating the effects of the other QTLs (Fig. 1e) and of unequal *P. polionotus* ancestry across individuals (Extended Data Fig. 2), both of which can influence burrow length. This approach resulted in a congenic mouse line with *P. polionotus* ancestry from chr4: 89.3 – 101.3 Mb on a *P. maniculatus* genomic background (Fig. 2a-c; Extended Data Fig. 3); the introgressed region is a near-perfect match to the 95% Bayes credible interval of the QTL-chr4 (Fig. 2c). By interbreeding mice heterozygous for this chr4 region, we generated sibling congenic mice representing the three QTL-chr4 genotypes: homozygous *P. polionotus* (*p*/*p*), heterozygous (*m*/*p*) and homozygous *P. maniculatus* (*m*/*m*) (Fig. 2d). Because we tested siblings with shared parents and shared home cages, our design minimizes environmental effects.

**Fig. 3.**
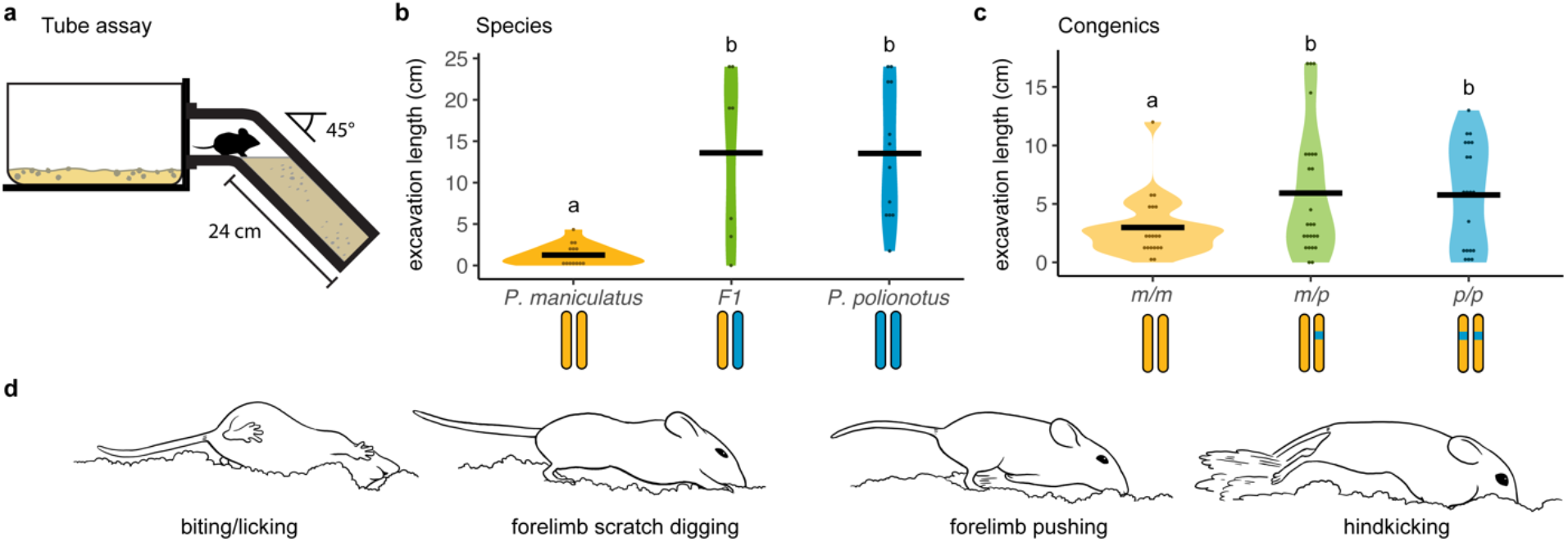
A novel tube assay to measure mouse burrowing behavior. (**a**) Diagram of the tube assay used for recording burrowing behaviors, including standard home cage attached to downward tilting plexiglass tube filled with sand. (**b**) Length of excavation in the tube assay for *P. maniculatus* (*n* = 18), *P. polionotus* (*n* = 12) and their F1 hybrids (*n* = 7), with black lines denoting mean excavation length by group. Letters represent statistically different groups (K-W test with Dunn’s test; *P* < 0.004). (**c**) Length of excavation for congenic mice of each genotype (*m/m* (*n* = 25), *m/p* (*n* = 26), *p/p* (*n* = 20)), with black lines denoting mean excavation length by genotype. Letters represent statistically different groups (Wilcoxon test; *P* < 0.05). (**d**) The four most frequently observed *Peromyscus* burrowing behaviors. Images drawn from video stills of a *P. polionotus* individual in the tube assay.

We next characterized the effects of QTL-chr4 by measuring the burrows dug by congenic siblings of all three genotypes in the large sand enclosures. We found that heterogyzous (*m*/*p*) congenic mice constructed significantly longer burrows than congenic *P. maniculatus* (*m*/*m*) mice (Fig. 2e), with average burrow lengths of 13.6 and 11.0 cm, respectively (two-sided t-test: *P* = 0.025). This result confirms that QTL-chr4, on its own, affects burrowing behavior: the *P. polionotus* allele increases burrow lengths by 2.6 cm on average in the congenic mice (a 23.6% increase).

Since genetic mapping was performed in backcross mice (which produce only *m*/*m* and *m*/*p* genotypes), the extent of dominance of QTL-chr4 was unknown. To investigate dominance, we also measured burrowing of *p*/*p* congenic mice and found they constructed burrows that were, on average, 13.3 cm long, significantly longer than the burrows constructed by *m*/*m* mice (two-sided t-test: *P* = 0.043), but indistinguishable from *m*/*p* mice (two-sided t-test: *P* > 0.05) (Fig. 2e). This result reveals the dominance of the QTL-chr4 and comports with the overall dominance of *P. polionotus* burrowing relative to *P. maniculatus*^11, 12^.

## Novel assay to directly measure digging behavior

We next explored how the *P. polionotus* allele at QTL-chr4 may affect digging behavior to increase burrow length. To this end, we designed a novel assay to capture individual components of burrowing behavior. In brief, the arena is comprised of a housing cage connected to a sand-filled tube with a transparent face, which facilitates video recording of mice as they burrow into the tube (Fig. 3a). The tube’s width (3.8 cm) and angle (45°) are intermediate to those typical of the parental species (*P. maniculatus* and *P. polionotus*), and tube length (24 cm) is sufficient to accommodate the downward-sloping portion of both species’ burrows^14^.

As a first step, we confirmed the tube assay recapitulates the species-specific behaviors by testing the parental species. After one hour in the assay, *P. polionotu*s dug significantly greater lengths (hereafter referred to as “excavation length”) than *P. maniculatus*, while F1 hybrids were statistically indistinguishable from *P. polionotus* but distinct from *P. maniculatus* (average excavation length: *P. polionotu*s = 13.5 cm, SD = 8.08; *P. maniculatus* = 1.25 cm, SD = 1.20; F1 hybrids = 13.6 cm, SD = 10.2) (Fig. 3b). These data confirm that the tube assay recapitulates the species-specific burrow lengths seen in nature^12, 15–17^ and observed in our sandbox assays (see Fig. 1a-c).

To test the specific effects of QTL-chr4 on burrowing, we evaluated congenic siblings of each genotype in the tube assay. We found QTL-chr4 produced a significant effect on excavation length: *m*/*p* and *p*/*p* mice removed significantly more sand than *m*/*m* mice, with the *P. polionotus* allele again showing dominance (average length dug in tube: *m*/*m* = 2.98 cm, SD = 2.46; *m*/*p* = 5.93 cm, SD = 5.45; *p*/*p* = 4.45 cm, SD = 4.45) (Fig. 3c). On average, mice harboring *P. polionotus* ancestry at QTL-chr4 dug more than those homozygous for the *P. maniculatus* allele by 2.88 cm in tube assays, comparable to our results in sandbox assays (2.6 cm).

In addition to measuring excavation length, our tube assay allows detailed characterization of the specific behavioral components of burrowing. Through observation of both parental species in the tube, we found mice moved and interacted with the substrate using both their limbs and their heads. We cataloged a repertoire of six distinct digging behaviors expressed by both species (Table S1); no species-specific behaviors were identified. Four of the six motor behaviors were observed frequently (Fig. 3d; >5 average observations per species): biting/licking packed sand, digging packed sand with the forelimbs, pushing loosened sand with the forelimbs, and kicking loosened sand with the hindlimbs (hereafter “hindkicks”).

## Species-specific differences in digging behavior

We first investigated how *P. polionotus* burrowing behavior may differ from that of *P. maniculatus*. We focused on two general behavioral mechanisms: (1) behavioral metrics associated with the drive to dig (e.g., temporal aspects of digging) and (2) traits associated with use of motor patterns (Fig. 4). First, in the parental species, we found that *P. polionotus* had both a shorter latency to enter the tube (K-W test, chi-squared = 16.5, df = 2, *P* = 3 × 10^−4^; Extended Data Fig. 4c) and a shorter latency to start digging in the tube (K-W test, chi-squared = 16.0, df = 2, *P* = 3 × 10^−4^) than *P. maniculatus* (Fig. 4a). *P. polionotus* also showed greater engagement, spending more time inside the tube (ANOVA, *P* = 3 × 10^−3^) (Fig. 4c) and more of that time digging (ANOVA, *P* = 9 × 10^−4^) (Fig. 4e) than *P. maniculatus*. Of these behaviors, only time spent digging showed a pattern of *polionotus*-dominance in the F1 hybrids (Fig. 4e), suggesting this trait may be the most consequential for burrow length. Together, these results are consistent with *P. polionotus* having greater drive to burrow than *P. maniculatus*, which may represent an innate difference in internal state.

**Fig. 4.**
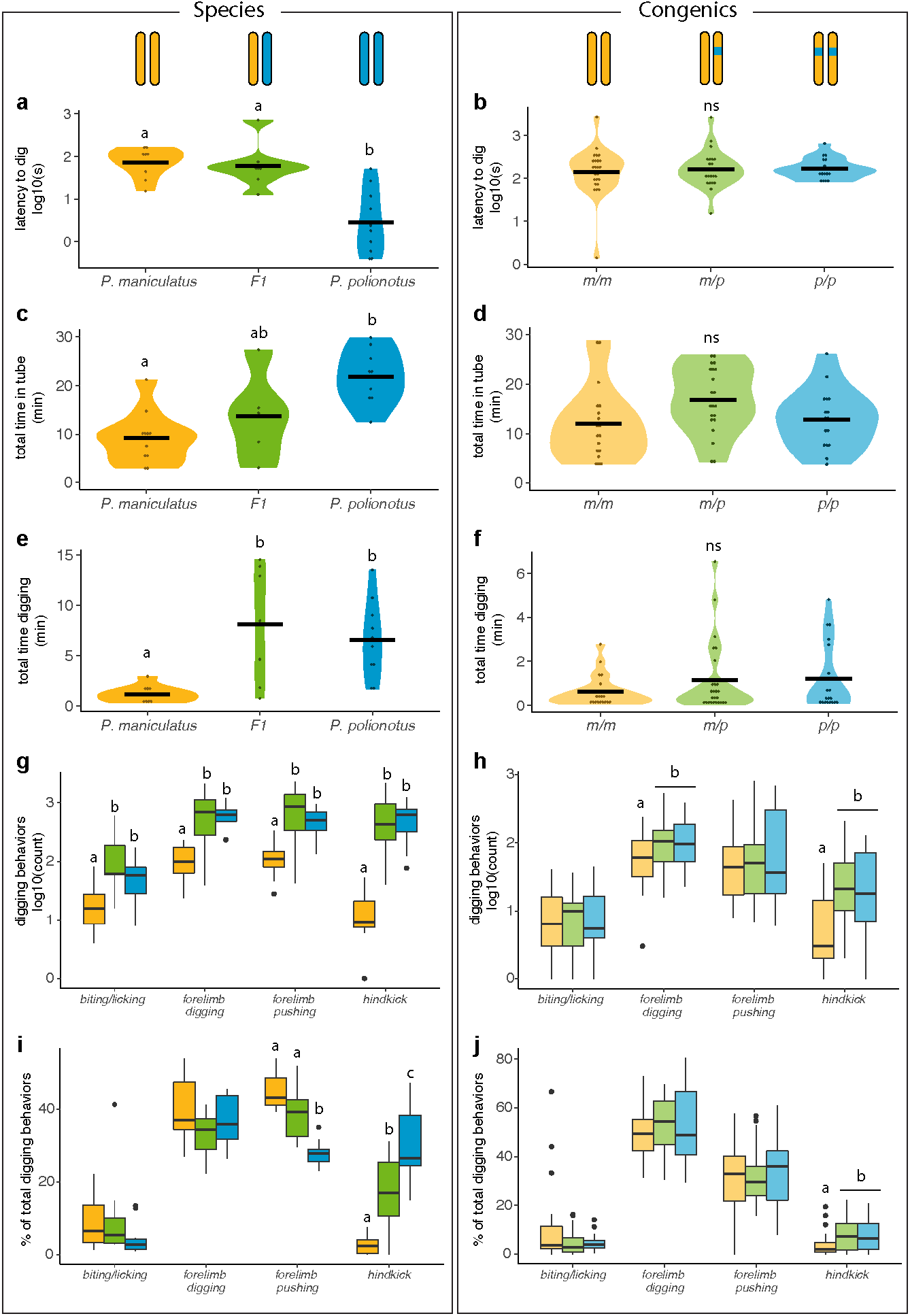
Effects of QTL-chr4 on distinct components of burrowing behavior. Measurements from the tube assay shown for the two species and their F1 hybrids (left column) and the three genotypes in the congenic line (right column). (**a-b**) Time (sec) until the first digging behavior observed for each mouse. (**c-d**) Total amount of time (min) each mouse spent inside the tube. (**e- f**) Total amount of time (min) each mouse spent digging in the tube. (**g-h**) Count of each digging behavior, shown on the log scale, per trial. (**i-j**) Proportional use of each digging behavior, calculated as counts of the specific digging behavior divided by counts of all behaviors. For (a-f), black lines denote mean by group; dots represent individual mice. In (g-j), boxplots show median (black line), with interquartile range; whiskers show minimum and maximum. Dots represent outliers. Letters represent statistically different groups (left side, K-W test with Dunn’s test, *P* < 0.05; right side, Wilcoxon test, *P* < 0.05). Species sample sizes: *P. polionotus* (*n* = 10), *P. maniculatus* (*n* = 11), F1 hybrids (*n* = 7). Congenic sample sizes: *m/m* (*n* = 25), *m/p* (*n* = 26), *p/p* (*n* = 19).

Second, we focused on differences in motor pattern, testing the hypothesis that *P. polionotus* digs longer burrows because of differences in the usage (i.e., count, proportion or latency) of the specific digging behaviors it performs. We first note that because *P. polionotus* spends more time digging, this resulted in an increase in the counts of the four common digging behaviors (see Fig. 3d) compared to *P. maniculatus* (K-W test followed by Wilcoxon with BH adjustment: biting/licking, *P* = 6 × 10^−3^; forelimb digging, *P* = 6 × 10^−4^; forelimb pushing, *P* = 2 × 10^−3^; hindkick, *P* = 4 × 10^−4^) (Fig. 4g). However, the proportional usage of specific digging behaviors performed (i.e., count of a specific behavior divided by total counts) also differed between the two species. Specifically, *P. polionotus* showed greater proportion of hindkicks (Wilcoxon, W = 0, *P* = 1.2 × 10^−4^; Fig. 4i), about ten times that of *P. maniculatus* (*P. polionotus*: mean = 30.4%, SD = 10.2%; *P. maniculatus*: mean = 2.75%, SD = 2.54%; Cohen’s *ds* =3.8). Moreover, *P. polionotu*s had a shorter latency to use hindkicks after initiating digging than *P. maniculatus* (Extended Data Fig. 5a; Wilcoxon test, W = 76, *P* = 0.0005). In sum, we found that *P. polionotus* construct longer burrows due to a combination of increased drive to dig and increased usage of a specific digging behavior, namely hindkicks.

**Fig. 5.**
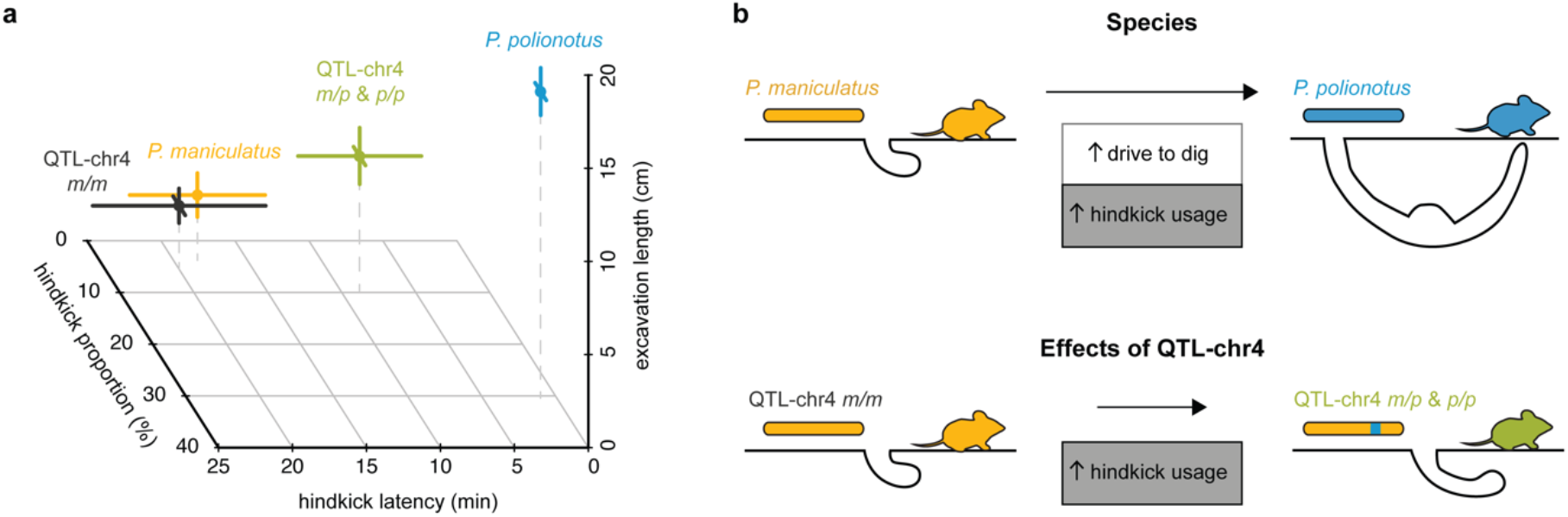
Hindkick-specific effects of QTL-chr4. (**a**) Hindkick proportion (proportion of digging behaviors as hindkicks), hindkick latency (time to first hindkick normalized to digging onset) and excavation length (cm dug in the tube assay) for parental species (*P. polionotus*, blue, *n* = 10; *P. maniculatus*, yellow, *n* = 8) and congenic mouse genotypes (*m/p* & *p/p*, green, *n* = 36; *m/m*, black, *n* = 19). QTL-chr4 allele explains 22% and 56% of the differences in hindkick proportion and latency, respectively, between the parental species. (**b**) A simplified model showing the evolution of burrow length (not to scale). Increases in both drive to dig (white) and hindkick usage (grey) contribute to the longer burrows of *P. polionotus* relative to *P. maniculatus* (not to scale). QTL-chr4 has a specific effect on hindkick usage, not drive to dig.

## Effect of QTL-chr4 locus on digging behavior

We next investigated the effects of QTL-chr4 locus on these two mechanisms – the drive to dig and the usage of digging behaviors – by scoring behavior of congenic mice of all three genotypes in the tube assay. We found that the QTL did not have significant effects on temporal aspects associated with digging: the latency to dig (Fig. 4b), total time spent in the tube (Fig. 4d), or the total time spent digging (Fig. 4f), suggesting that QTL-chr4 does not influence drive to dig (Wilcoxon tests, *P* > 0.05). By contrast, we found a significant difference in the usage of digging behaviors among genotypes. While genotypes did not differ in the usage of biting/licking or forelimb pushing behavior (Wilcoxon tests, *P* > 0.05), we did see a difference in the count of forelimb digs (Wilcoxon, W = 389, *P* = 0.04) as well as hindkicks (W = 374.5, *P* = 0.03) (Fig. 4h). However, when we controlled for the total amount of digging behaviors performed (i.e., total counts), we found that the proportion of only hindkicks differed among genotypes (Wilcoxon test, W = 386, *P* = 0.04; Fig. 4j). Specifically, congenic mice carrying at least one *polionotus* allele (*m/p* and *p/p* genotypes) had a greater proportion of hindkicks (mean = 7.57%, SD = 6.45%) compared to *m*/*m* mice (mean = 4.17%, SD = 5.21%; Cohen’s *ds* = 0.56; Fig. 4j), with QTL-chr4 increasing the proportion by 22% of the difference observed between the parental species (Fig. 5a). Despite no observed difference in the latency to start digging among genotypes, QTL-chr4 did affect the latency of hindkicking (Extended Data Fig. 5b), explaining 56% of the difference observed between the parental species (Fig. 5a). The differences in hindkicking between genotypes appeared early and persisted (Extended Data Fig. 5c). Notably, hindkicking behavior began when burrows were still short (Extended Data Fig. 5d; Wilcoxon, *P* > 0.05), demonstrating that hindkicking is not used to eject soil exclusively from long burrows. In sum, we found that while *P. polionotus* build long burrows through a combination of both increased drive to dig and increased hindkick usage, QTL-chr4 has a measurable and specific effect on only hindkick usage (Fig. 5b).

## Discussion

Here we identify, isolate, and then test how a single genomic locus contributes to the elaboration of a complex natural behavior. Functional characterization of *natural genetic variation* remains rare in the field of behavioral genetics (but see ^18–20)^, especially in mammals. By isolating a 12-Mb region (<0.5% of the 2.5 Gb genome) from the long-burrow species onto a consistent genetic background of the short-burrow species, we demonstrate that this locus alone acts dominantly to increase burrow length by >20%, providing both strong causal evidence that this region contributes to burrow evolution and allowing us to parse its specific effects on a complex, multicomponent behavior.

The development of a novel behavioral assay, which captured both the temporal dynamics and discrete motor behaviors of deer mouse burrowing behavior – that occurs underground and at night – allowed for a unique lens into the evolution of different burrow architectures. Interestingly, we did not discover any new motor patterns for digging in the long-burrowing *P. polionotus*, suggesting that evolution acted on only existing behavioral components, contrasting with other examples of behavioral evolution, such as novel motor patterns observed in sea slug swimming^21^ and snake climbing behavior^22^ as well as new courtship components of avian displays^23^ and cricket sound^24^. Instead, two behavioral mechanisms contribute to the construction of long burrows. First, *P. polionotus* show an increased drive to dig compared to *P. maniculatus*, which results in more time spent digging and higher counts of the four most frequent digging behaviors. Second, the species differ in action selection for specific motor patterns: hindkicks represent a greater proportion of digging behavior in *P. polionotus* than *P. maniculatus*. These results suggest that changes at both a higher level of internal state (i.e., drive, or colloquially, “motivation”) and a lower level (i.e., use of specific motor patterns) of behavioral organization contribute to the evolution of burrowing between these two species, although the relative contribution of each mechanism is unclear.

Moreover, these ostensible multi-level differences between parental species allows for the possibility that individual loci could act at multiple, or only specific, levels of this hierarchy. Previous studies have shown that behaviors can be disentangled into genetically discrete modules at “high” phenotypic levels, for example, different physical aspects of burrow architecture (entrance versus escape tunnels)^12^, features of fish schooling behavior (tendency versus position)^25^, types of sound production (ultrasonic versus audible)^26^ or aspects of parenting behavior (nest building versus pup retrieval)^27^. We investigated this question by testing how a single genetic region—isolated in a congenic line—contributes to burrow length differences at a fine-scale behavioral level. Surprisingly, this region affected only usage of specific motor patterns, while showing no effects on measures of drive to dig (e.g., latency to dig and time spent digging). This is consistent with it acting specifically at a lower level of the hierarchy, and providing a rare example of a genetic change that influences action selection for motor patterns. Further, it demonstrates that the two behavioral mechanisms that lead *P. polionotus* to dig long burrows—increased drive to burrow paired with distinct motor pattern use—are genetically discrete.

Most strikingly, the locus increased usage of hindlimb kicking. Hindkicks have been described as powerful “blasts” that can propel soil >60 cm rearward^28^ (Extended Data Video 1). Because hindkicks involve synchronized (compared to alternating, as in forelimb digging) limb movement, they resemble bounding and galloping gaits (compared to walking/trotting gaits). In neurobiology, gait transitions have been a model for studying central pattern generators, the local circuits in the spinal cord that generate locomotion^29^. At the circuit level, different neuronal ensembles are being recruited to carry out different motor patterns, and thus coordinating interneurons are an attractive candidate substrate for the evolution of motor pattern selection^9, 29, 30^. At the genetic level, gait variation in domesticated show horses has been linked to *Dmrt3*, a gene that plays a role in interneuron specification in the spinal cord^31^. While this gene is not on the same chromosome as QTL-chr4, future work to further characterize the QTL-chr4 locus, which contributes to *natural* variation in motor pattern selection, will provide novel insights—and a unique genetic approach—into the mechanistic basis of behavioral evolution.

Fifty years ago, Tinbergen and two colleagues were awarded a Nobel Prize for their studies on the evolution of behavioral complexity. Here we show that individual genetic loci can have surprisingly specific effects on behavior, providing empirical support at the *genetic level* for the prize-winning idea that behavior can be organized hierarchically. This structure, in turn, provides evolution the opportunity to make fine-tuned behavioral changes, thereby facilitating behavioral adaptation of animals to their local environments.

---

**Extended Data Fig. 1.**
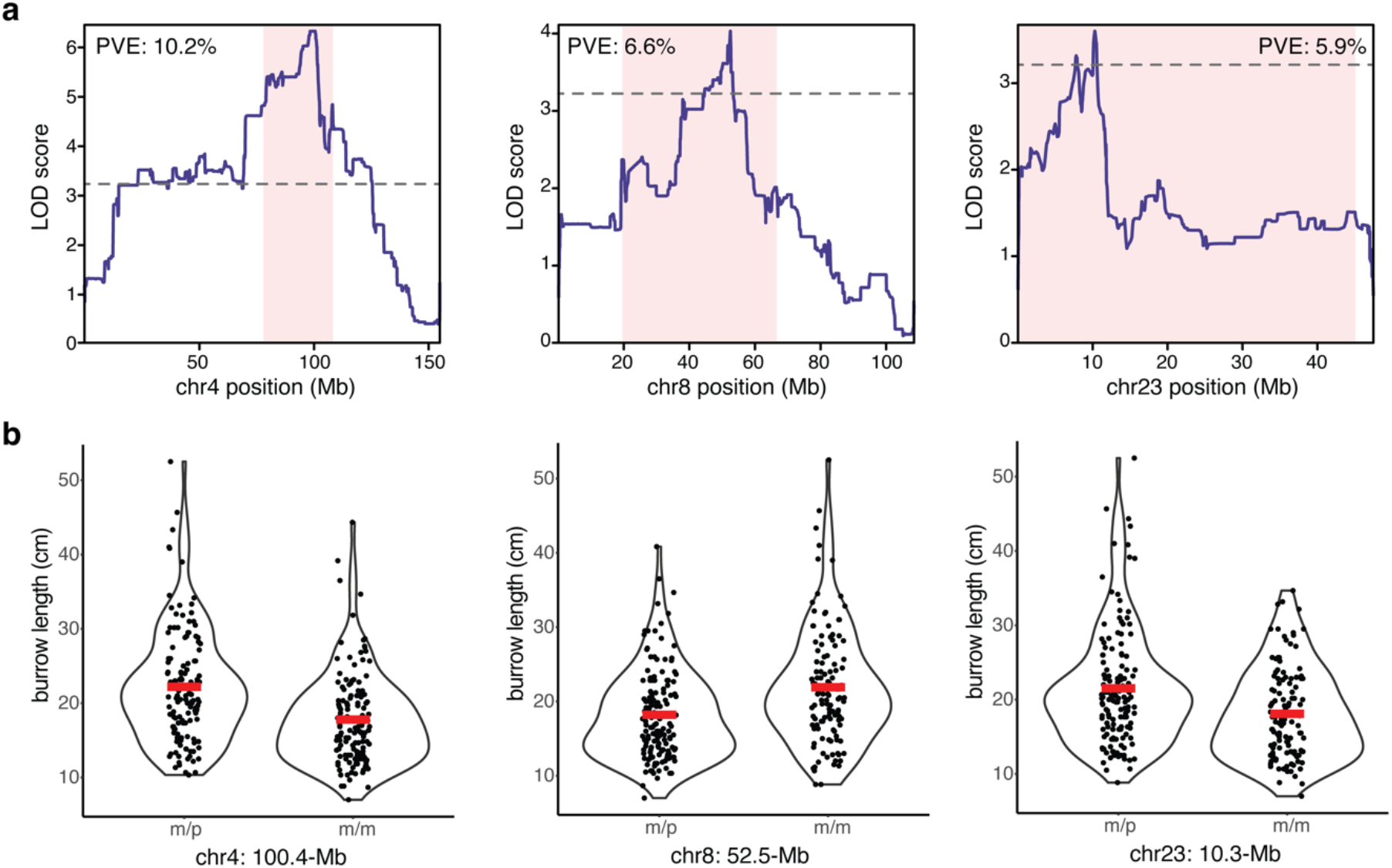
Burrow length QTL effects in BC1 hybrids. (**a**) Association between burrow length and genotype for the three significant burrow length QTL on chromosomes 4, 8 and 23. Shaded regions (pink) show 95% Bayes credible interval; dashed line indicates genome-wide significance threshold. PVE = percent variance explained. (**b**) Effect sizes for the three QTLs. BC1 hybrids are grouped by genotype at the peak marker for each QTL. Dots represent individuals; mean burrow lengths by genotype are shown (red lines). *m*/*p* = heterozygous *P. maniculatus*/*P. polionotus*; *m*/*m* = homozygous *P. maniculatus*. Sample size: chr4: *m/p* (*n* = 132), *m/m* (*n* = 139); chr8: *m/p* (*n* = 151), *m/m* (*n* = 118); chr23: *m/p* (*n* = 140), *m/m* (*n* = 124).

**Extended Data Fig. 2.**
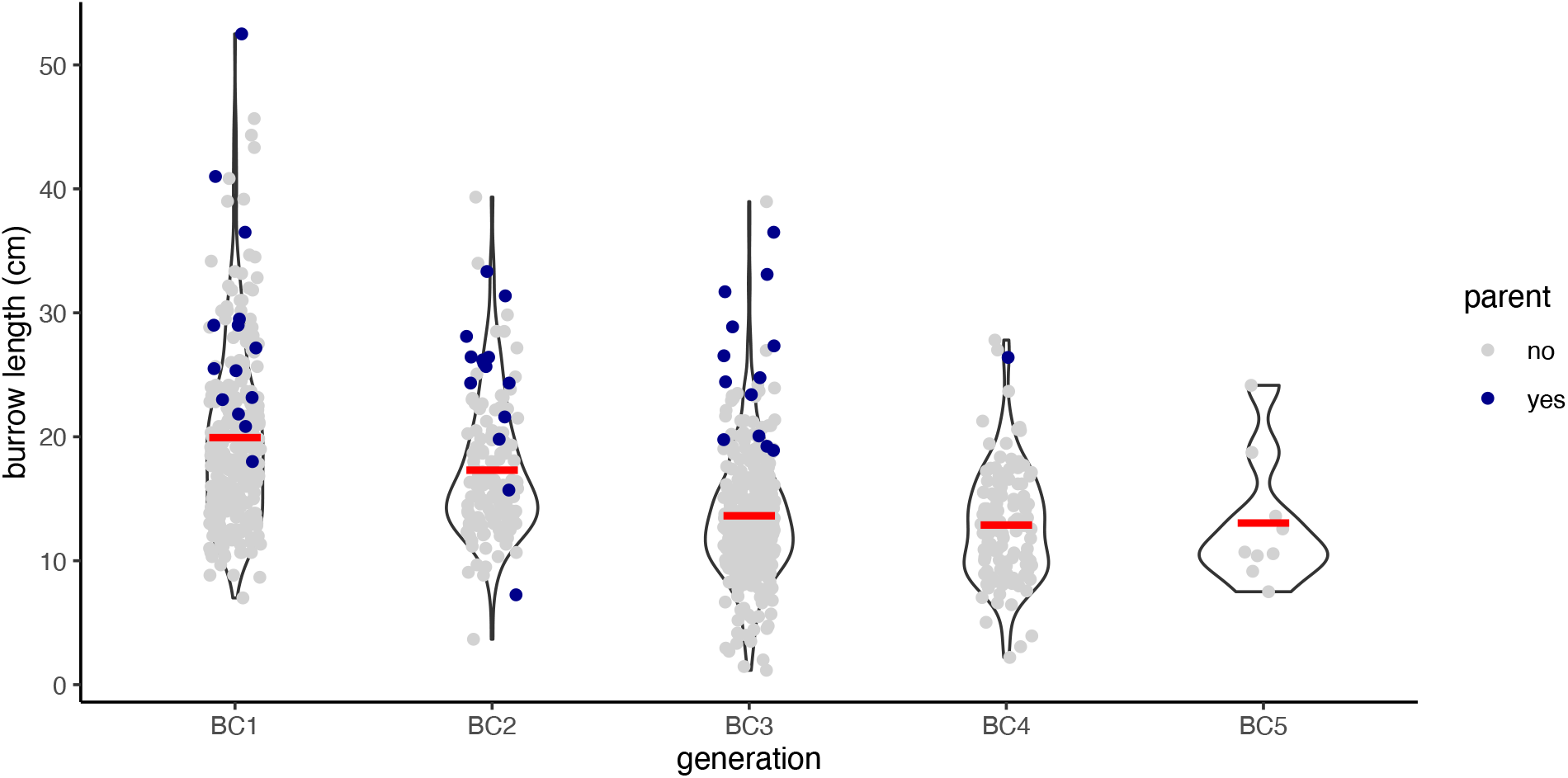
Burrow length by advanced backcross generation. Burrow lengths are shown by advanced backcross generation (BC1-BC5), with mean burrow length per generation denoted by red lines. Dots represent individual BC hybrids; mice selected as a parent of the next BC generation are highlighted (blue dots). Sample size: BC1 (*n* = 271); BC2 (*n* = 152); BC3 (*n* = 330); BC4 (*n* = 130); BC5 (*n* = 9).

**Extended Data Fig. 3.**
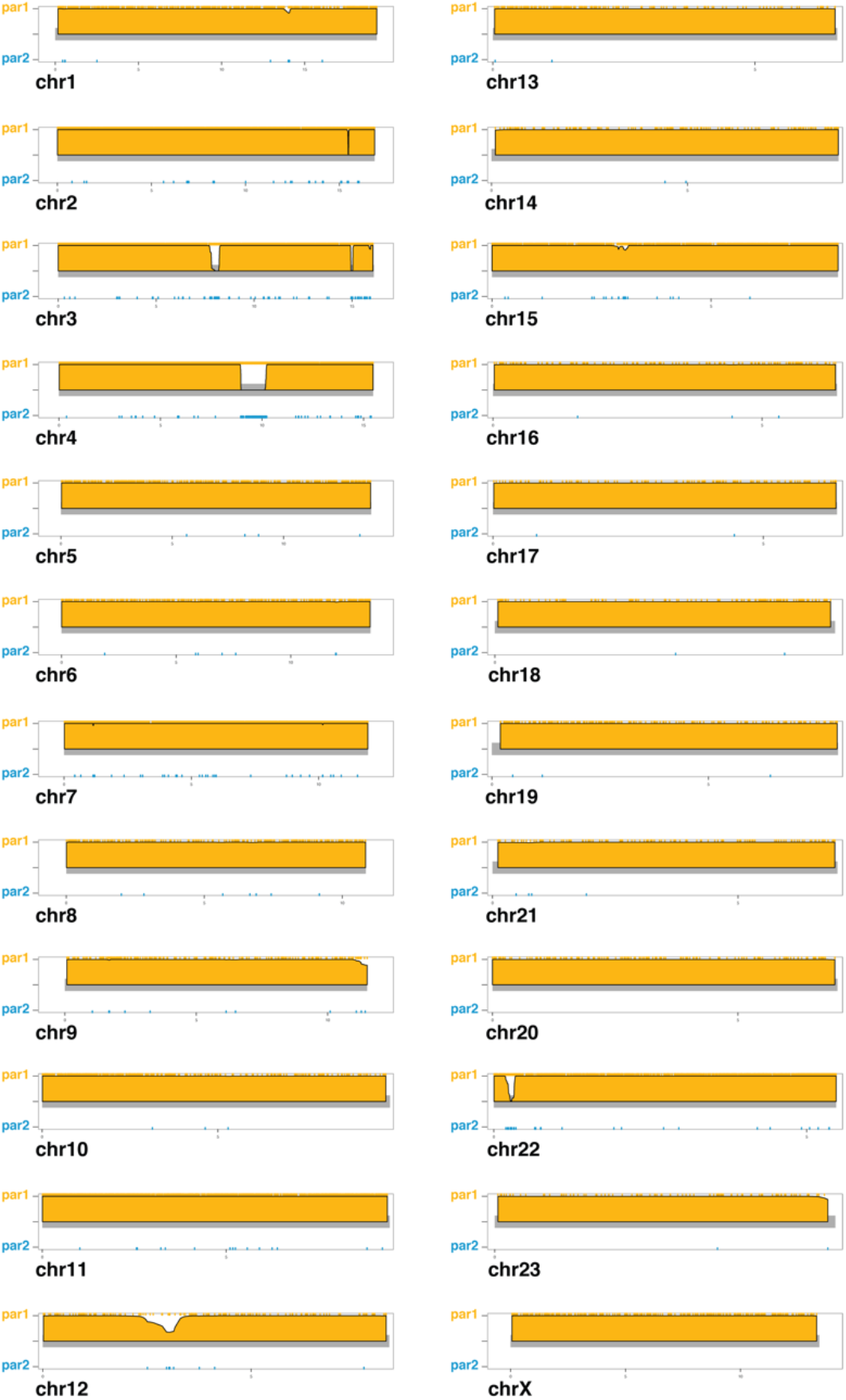
Genome-wide ancestry for QTL-chr4 congenic mouse line founder. Ancestry probabilities for each chromosome for the founder mouse of the QTL-chr4 congenic mouse line. Tick marks represent ancestry informative SNPs (yellow: *P. maniculatus*; blue: *P. polionotus*). Y-axis shows ancestry probabilities (par1: *P. maniculatus*, yellow; par2: *P. polionotus*, blue) from the hidden Markov model (see Methods). Heterozygous regions (equal probabilities of both ancestries) are shown as gray lines (e.g., chromosome 4).

**Extended Data Fig. 4.**
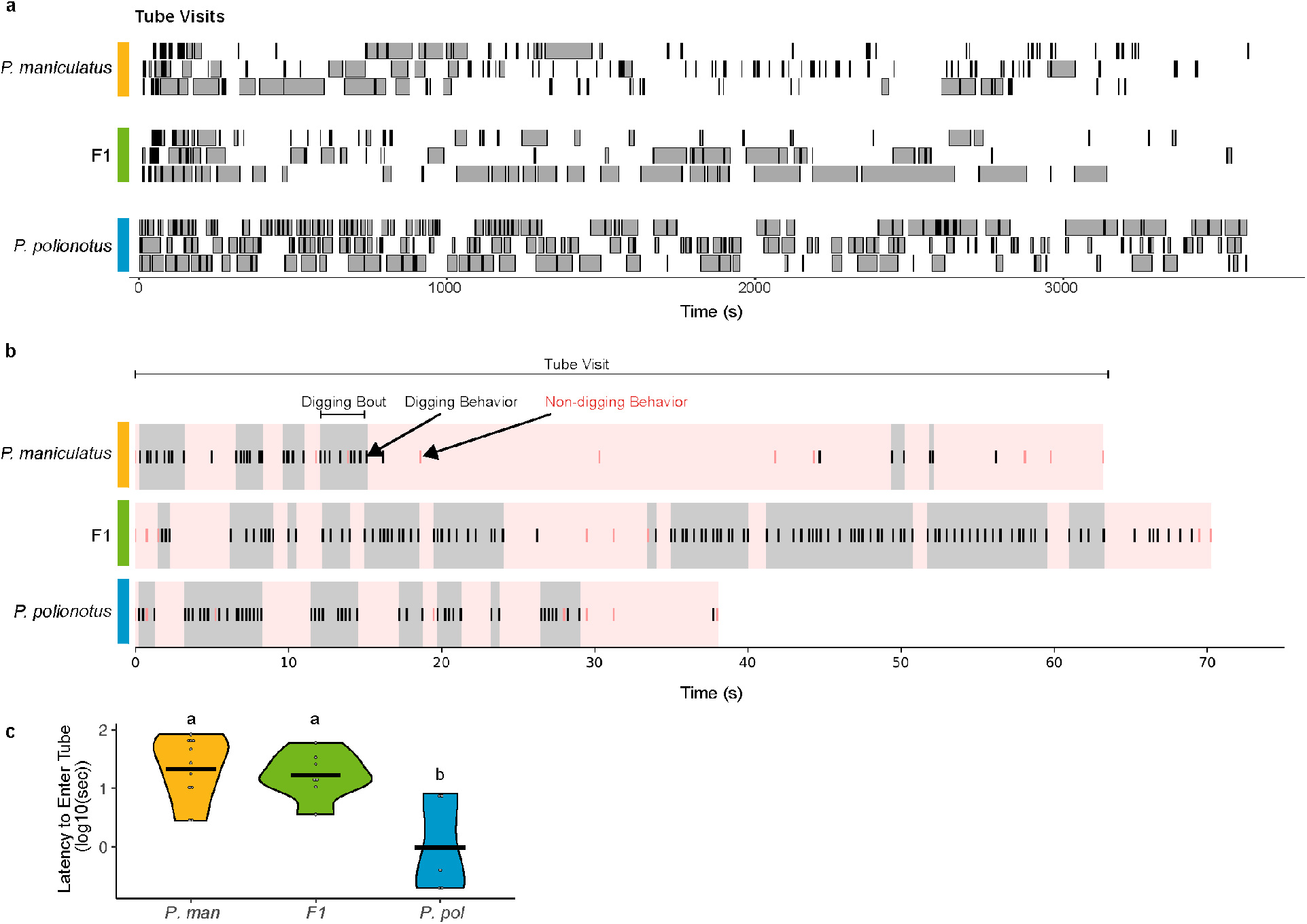
Examples of tube visit and digging bouts. (**a**) Representative 1 h trials for *P. maniculatus*, F1 hybrids, and *P. polionotus* (*n* = 3 mice per group). Visit durations (defined as the time from which a mouse entered a tube to when it exited) are plotted as rectangles. Visits during which the animal performed a digging behavior are filled in gray and visits during which the animal did not are filled in black. (**b**) A single tube visit, containing the median number of digging bouts, for each group is shown. The total duration of the tube visit is highlighted (pink). Digging bouts (defined as when digging behaviors occurred in succession with <1 s between digging behaviors) during tube visits are highlighted (gray). Tick marks for individual behaviors: digging behaviors (black) and non-digging behaviors (dark pink). (**c**) The latency to enter the tube for each group. Bars indicate group means; letters indicate statistically different groups (K-W test with Dunn’s test, *P* < 0.05).

**Extended Data Fig. 5.**
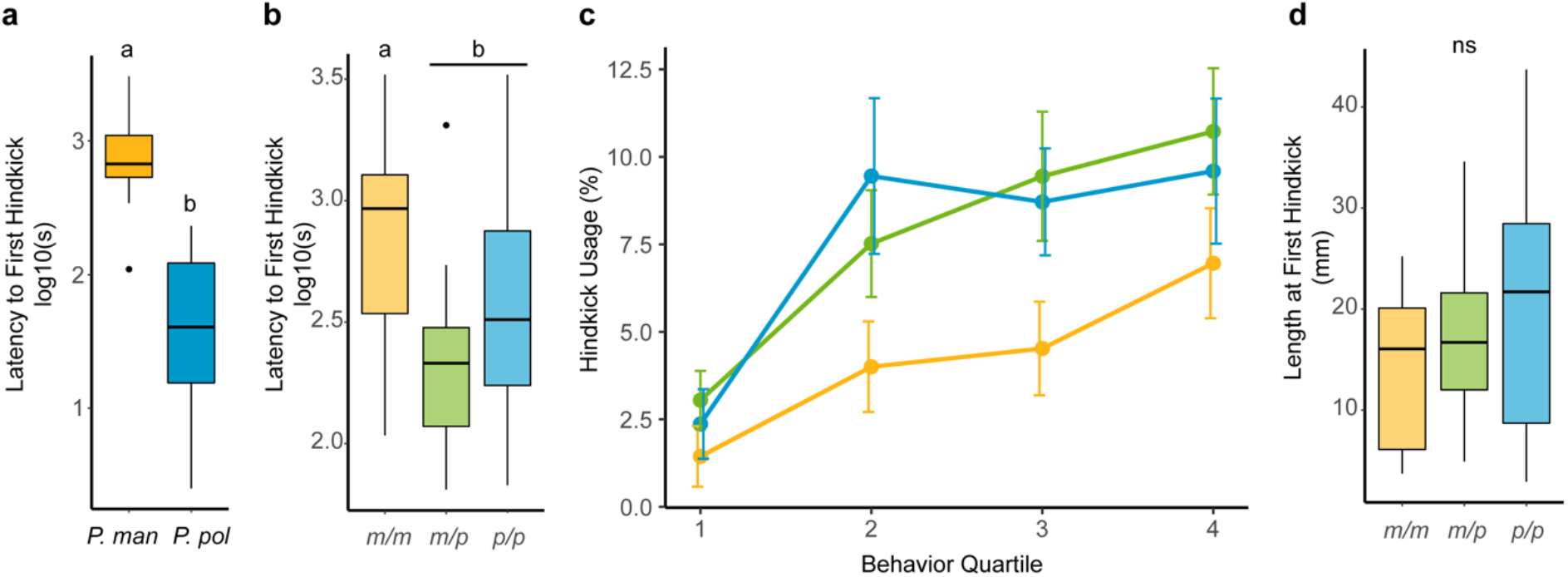
Effect of QTL-chr4 on latency to perform hindlimb kicking. (**a**) Latency to perform hindkicks (time to first hindkick after the onset of digging) in parental species (Wilcoxon test, *P* = 0.0005). Species sample sizes: *P. polionotus* (*n* = 10) and *P. maniculatus* (*n* = 8). (**b**) Latency to perform hindkicks among congenic genotypes (Wilcoxon test, *P* = 0.002). Sample sizes: *m/m* (*n* = 24), *m/p* (*n* = 26), *p/p* (*n* = 20). (**c**) The percent hindkicks of all digging behaviors for each quarter of each trial are shown for the congenic mice. Mice carrying the *P. polionotus* allele (green, blue) show increased hindkick usage in the second quarter and third quarter compared to those homozygous for the *P. maniculatus* allele (yellow) (*P* = 0.03, linear mixed-effects model with genotype as a factor and mouse ID as a random effect). Sample sizes same as (**b**). Dots represent means; bars represent standard error of the mean (SEM). (**d**) Length of burrow at which mice initiate hindkicks among congenic genotypes (Wilcoxon test, *P* > 0.05). Sample sizes: *m/m* (*n* = 18), *m/p* (*n* = 20), *p/p* (*n* = 16). For (**a**, **b**, and **d**), boxplots show median (black line) with interquartile range; whiskers show minimum and maximum. Dots represent outliers. Letters indicate statistically different groups.

## Methods

### Animal husbandry

Outbred colonies of *Peromyscus maniculatus bairdii* and *Peromyscus polionotus subgriseus* were originally derived from the Peromyscus Genetic Stock Center (University of South Carolina, Columbia, SC, USA). We performed all experiments using mice raised in controlled laboratory conditions. Mice were housed in ventilated polysulfone cages (19.7 x 30.5 x 16.5 cm, Allentown, New Jersey, USA) in same sex groups. Mice were maintained at 21C on a 16:8 h light:dark cycle and had *ad libitum* access to water and irradiated food (LabDiet Prolab Isopro RMH 3000 5P75, LabDiet, St. Louis, MO). We provided all housing cages with corncob bedding and polycarbonate translucent red hut.

We performed genetic crosses between *P. maniculatus* and *P. polionotus* to produce both first-generation (F1) and backcross (BC) hybrids, as well as the congenic mouse line (see “*Genetic mapping*” and “*Congenic mouse line*”, respectively, below for genetic cross details). All mice were maintained under the same conditions: mice used to start the cross (“founders”), their F1 hybrids, *P. maniculatus* “breeders” used to generate the BC hybrids, the BC hybrids, and the congenic mice. All procedures were approved by the Harvard University Institutional Animal Care and Use Committee.

### Burrowing assay: large sandboxes

We measured mouse burrow construction using large indoor PVC enclosures (1.2 x 1.5 x 1.1 m) filled with 700 kg of hydrated hard-packed premium play sand (Quickrete, Atlanta, GA) as previously described^12^. In brief, before introducing mice to the enclosures, we shaped the sand into a hill with three sections: a flat lower section, a middle hill angled at approximately 60 degrees from horizontal, and a flat upper section. In the enclosure, we provided each mouse with 5g of mouse chow (LabDiet Prolab Isopro RMH 3000 5P75), one cotton nestlet (Ancare, Bellmore, NY, USA), and one water bottle. For each trial, we introduced a single mouse to an enclosure at the start of the dark cycle and then removed the mouse approximately 45 h later. We recorded the locations and measured the lengths of all sand “excavations”. For excavations >8cm in length, we injected them with polyurethane filling foam (Hilti Corp., Schaan Liechtenstein) to make a physical cast. After the foam dried, we measured the total length of the excavation directly from the cast. We tested each mouse three times, consecutively, when mice were between 60 and 100 days of age and assigned mice to new enclosures at random for each trial.

We included both males and females, since burrowing does not differ between the sexes^12^. We summarized burrowing for each mouse using the average *burrow length* over the three trials, where *burrow length* is the length of the longest excavation created by the mouse during the trial.

### Genetic mapping

To describe the genetic architecture of burrow length measured in the large sandboxes, we first re-analyzed mice from a previous single-generation backcross^12^ (BC1) and then further fine-mapped within the identified genomic regions using a five-generational advanced backcross (BC1-BC5); these mapping approaches are described below. All mice used in the genetic crosses were sequenced using a ddRAD-sequencing approach^32^ (see *Library preparation and sequencing*) and genotyped using a hidden-Markov model approach^33^ (see *Genotyping backcross hybrids*).

#### Library preparation and sequencing

To determine individual genotypes, we used a ddRAD-sequencing approach^32^. In brief, we extracted DNA from tail, ear or liver tissue using an automated phenol-chloroform platform (Autogen, Holliston MA). For each individual, we prepared a library of restriction cut-site-specific genome-wide fragments using the enzymes EcoR1 and MspI (New England Biolabs, Ipswich MA), custom adapters (Integrated DNA Technologies, Coralville IA), and T4 ligase (New England Biolabs, Ipswich MA). We size-selected these libraries using a Pippin Prep (Sage Science, Beverly MA) and then amplified and indexed them via PCR (Phusion, New England Biolabs, Ipswich MA). At each stage, we purified fragments using Ampure XP beads (Beckman Coulter Genomics, Danvers MA) and assessed library quality using a Bioanalyzer (Agilent Technologies, Santa Clara, CA). We then sequenced these libraries (paired-end 2x150bp) on an Illumina HiSeq (San Diego CA). Note that the *P. maniculatus* and *P. polionotus* cross founders, F1 hybrids, *P. maniculatus* breeders, and BC hybrids were all included in the ddRAD-sequencing pipeline.

#### Genotyping backcross hybrids

To determine ancestry-informative genotypes, we first de-multiplexed the fastq files and then mapped the sequence reads to the *P. maniculatus* reference genome (Pman2.1.3) using BWA-MEM. Next, we merged sam files using *MergeSamFiles* (Picard) into a single indexed and sorted bam file per sample. We validated bam files using *ValidateSamFiles* (Picard).

To identify ancestry-informative SNPs between the *P. polionotus* and *P. maniculatus* cross founders, we first performed joint genotyping using GATK (v3.8). Specifically, we created per-sample gvcfs using *HaplotypeCaller*, with the default heterozygosity prior set to 0.001. We performed multi-sample joint genotyping using *GenotypeGVCFs*, resulting in one vcf file for each chromosome. SNPs were selected and filtered using *SelectVariants* based on GATK hard-filtering best practices recommendations: SNPs with QD<2.0, FS>60.0, MQ<40.0, MQRankSum< -12.5, or ReadPosRankSum< -8.0 were removed. We next selected SNPs with different homozygous genotypes between the *P. maniculatus* and *P. polionotus* founders (i.e., homozygous for the reference allele in one founder, homozygous for the alternate allele in the other founder), resulting in 431,640 SNPs. To ensure that these SNPs were fixed for the same allele in all of the *P. maniculatus* breeders used in the backcross (including the advanced BC, see below), we used *VariantFiltration* to remove any SNP that had a read depth greater than 1 and was not homozygous for the *P. maniculatus* founder’s allele in all of the *P. maniculatus* breeders, which removed ∼7.5% of the SNPs, resulting in a total of 398,987 SNPs. Finally, we used sequencing from 11 F1 hybrids to filter SNPs that were not heterozygous in the F1 hybrids. Due to the low coverage of our sequencing data (and thus low confidence in calling heterozygous sites), we pooled the F1 data to obtain overall allele frequencies. Then based on a binomial distribution where *n* is the total number of reads at that SNP across all F1 hybrids and *p* is 0.5, we filtered SNPs in which the alternate allele frequency was in the bottom or top 10^th^ percentile of the binomial distribution, thus removing SNPs whose allele frequency deviated from the expectation of 0.5 in the pooled F1 data. In the end, our approach resulted in a total of 160,624 high-confidence ancestry-informative SNPs.

We used the hidden Markov model implemented in the Multiplexed Shotgun Genotyping (MSG) pipeline^33^ to assign ancestry in the BC hybrids. In brief, we used *mpileup* (samtools) to extract the set of 160,624 ancestry-informative SNPs from the mapped bam files of BC hybrids, requiring that SNPs were at least 500 bp apart to ensure they were from independent reads, resulting in 98,452 SNPs. If a sample had fewer than 100 informative markers for a given chromosome, we excluded that chromosome since ancestry determination fails at low SNP density. Then, we ran the fit-HMM step of MSG (fit-hmm.R) on the filtered mpileups with the following settings: deltapar1=0.1, deltapar2=0.1, rfac=1; theta=1; one_site_per_contig=1; recRate=25; and generation-specific priors [autosomes (MM, MP, PP), male X-chr (MM, PP); BC1: autosomes=(0.5,0.5,0.0), X-chr=(0.75,0.25); BC2: autosomes=(0.75,0.25,0), X- chr=(0.875,0.125); BC3: autosomes=(0.875,0.125,0), X-chr=(0.9375,0.0625); BC4: autosomes=(0.9375,0.0625,0), X-chr=(0.969,0.031); BC5: autosomes=(0.969,0.031,0), X- chr=(0.9844,0.0156)].

#### QTL mapping

*First-generation backcross*: We first characterized the genetic architecture of burrow length using re-analyzed (previously unreported) burrow traits and newly generated sequencing data (described above) from BC1 hybrids first described by Weber and colleagues^12^. Specifically, this cross was started with a single *P. polionotus* male and a *P. maniculatus* female that yielded 13 F1-hybrids (male and female), which were then backcrossed to *P. maniculatus* breeders *(n* = 13) to produce 271 BC1 hybrids. A backcross design was used to maximize mapping power^34^, as F1 hybrids have similar burrow lengths to *P. polionotus* (i.e., long burrows are dominant).

To identify regions of the genome that contain mutation(s) contributing to burrow-length variation, we performed QTL mapping of the BC1 hybrids using r/qtl2^35^. We combined the genotype probabilities (from the hidden-Markov model) across the BC1 hybrids using combine.py (https://github.com/JaneliaSciComp/msg/), which interpolates missing genotypes and prepared the output data for r/qtl2 using pull_thin (adapted from https://github.com/dstern/pull_thin) with the following settings (diffac=0; chroms=all; cross=bc). We then created an r/qtl2 cross object using an adapted version of read.cross.msg.1.5.R (https://github.com/dstern/read_cross_msg/). We mapped log(*burrow length*) such that the phenotype was normally distributed, since *burrow length* data across BC1 hybrids is right-skewed. QTL mapping was performed using *scan1* (r/qtl2) with Haley-Knott regression. To identify the LOD-significance threshold, we performed permutation tests using *scan1perm* (r/qtl2) with 1,000 permutations for the autosomes, and 17,144 permutations for the X-chromosome (determined from the *scan1perm* function using *perm_Xsp*). We then determined 95% Bayes credible intervals for the significant QTL peaks using *bayes_int* (r/qtl2).

To compare the QTL reported here with those previously described by Weber and colleagues^12^, we used a liftover bed file between the original linkage map and the current reference genome (Pman2.1.3); we found that the QTL on chromosomes 4 and 23 overlapped with the QTL described by Weber and colleagues on linkage groups 2 and 20, respectively.

*Advanced backcross*: To refine and narrow the boundaries of the major QTL located on chromosome 4 (“QTL-chr4”), we conducted an advanced backcross experiment. Specifically, we repeatedly backcrossed hybrid mice to *P. maniculatus*, using phenotypic selection at each generation. Starting with the BC1 generation, we selected hybrids (either males or females) according to their *burrow lengths* (Extended Data Fig. 2) and tunnel number (i.e., escape tunnels^12^), with a few exceptions to perpetuate family structure. Selected hybrids were then crossed to *P. maniculatus* breeders. We repeated this selection regime until we reached backcross generation 5 (BC5). The following number of mice were selected at each backcross generation to start the next generation: *n* = 14 (BC1), 14 (BC2), 13 (BC3), 1 (BC4). We measured burrow length in all offspring from each BC generation using the large sandbox assay (as described above), with three trials per mouse.

To narrow the QTL-chr4 region, we performed genetic mapping in r/qtl2^35^ using the genotypes as determined above. We first used the R-package *pedigree* to create a relatedness matrix based on the pedigree for all BC1 – BC5 hybrid mice; this matrix controls for the unequal relatedness and family structure within the advanced BC mapping population. To perform fine-mapping of QTL-chr4, we combined the genotype probabilities (from the hidden-Markov model) across the BC1-BC5 hybrids using combine.py (https://github.com/JaneliaSciComp/msg/) and prepared the output data for r/qtl2 using pull_thin (adapted from https://github.com/dstern/pull_thin) with the following settings (diffac=0; chroms=all; cross=bc). We then created an r/qtl2 cross object using an adapted version of read.cross.msg.1.5.R (https://github.com/dstern/read_cross_msg/). We performed genetic mapping in r/qtl2 using a linear mixed model, with relatedness as a random effect and log(*burrow length*) as the phenotype. The linear mixed model was implemented using *scan1* (r/qtl2), specifying the relatedness matrix from *pedigree* as a random effect. We then used *scan1perm* (r/qtl2) with 1,000 permutations to identify the LOD-significance threshold for chromosome 4. Then, using *bayes_int* (r/qtl2), we determined the 95% Bayes credible interval for QTL-chr4.

### Congenic mouse line

To characterize the specific effects of the QTL-chr4 region, we generated a congenic line by introgressing *P. polionotus* ancestry at the QTL-chr4 into a *P. maniculatus* genomic background. To create this congenic line, we backcrossed *P. polionotus* to *P. maniculatus* mice, using new founder mice from those described for QTL mapping. To determine ancestry in the QTL region, we designed two custom Taqman SNP genotyping assays (Life Technologies) to genotype the BC mice for diagnostic SNPs across the QTL-chr4 region (Table S2). These genotypes allowed us to determine *P. polionotus* versus *P. maniculatus* ancestry along the 12-Mb of QTL-chr4. We then selected BC hybrids containing *P. polionotus* ancestry across the QTL-chr4 region and crossed these selected mice with *P. maniculatus* breeders.

To identify the boundaries of *P. polionotus* ancestry on chromosome 4 as well as reduce *P. polionotus* ancestry across the rest of the genome, we performed low-coverage (∼0.5x coverage) whole-genome sequencing at BC generations 2 and 5. We prepared sequencing libraries using the NexteraXT library preparation kit and then sequenced them on an Illumina NextSeq mid-output flowcell at Janelia Research Campus, with 2x150bp paired-end reads. We mapped the sequence reads to the *P. maniculatus* genome (Pman2.1.3) with BWA-MEM, and sam files were sorted and indexed with *SortSam* (Picard) to create bam files. Duplicates were then marked with *MarkDuplicates* (Picard).

To determine ancestry across the genome, we identified a set of ancestry informative SNPs using high-coverage (∼15x coverage) whole-genome sequencing data from wild-caught *P. polionotus* (*n* = 15) and *P. maniculatus* (*n* = 17) mice (NCBI SRA PRJNA838595, PRJNA862503). Using a variant call file for the wild-caught individuals (see ^36^), we selected SNPs that were fixed as opposite homozygous genotypes between the *P. polionotus* and *P. maniculatus* mice, resulting in 39,990 – 388,630 SNPs per chromosome. This large set of SNPs was helpful for ancestry determination since the BC hybrids were sequenced at low coverage, with only a subset of SNPs covered in each mouse. We selected this set of ancestry-informative SNPs from the bam files of BC hybrids using *mpileup* (samtools) and then determined genome-wide ancestry using the fit-HMM step of MSG.

The whole-genome sequencing of BC hybrids revealed a BC family which harbored only four blocks of *P. polionotus* ancestry genome-wide, including an ancestry block covering QTL- chr4. This chromosome 4 ancestry block ended at chr4:101.3-Mb, which is only ∼300-kb from the 95% credible interval boundary for QTL-chr4. Through three additional backcross generations of mice from this family to *P. maniculatus*, we removed the additional *P. polionotus* ancestry blocks from this line. Further, a recombination breakpoint within the chromosome 4 ancestry block resulted in a BC hybrid with breakpoints at chr4:89.3-Mb and chr4:101.3-Mb, both breakpoints within ∼300-kb from the boundaries of the QTL-chr4 95% credible interval.

This recombination breakpoint at chr4:89.3-Mb was initially identified with the Taqman SNP assays and then verified with low-coverage whole-genome re-sequencing. Through generating offspring from this founder BC mouse, we obtained a congenic mouse line with *P. maniculatus* ancestry across the entire genome, except with *P. polionotus* ancestry at chr4: 89.3 – 101.3 and a few possible other short *P. polionotus* tracts (Extended Data Fig. 3). We then crossed seven pairs of mice from this line that were heterozygous (*P. polionotus* and *P. maniculatus* ancestry) at the QTL-chr4 region to obtain sibling offspring with the three possible genotypes for QTL-chr4: homozygous *P. polionotus* (*p*/*p*); heterozygous (*m*/*p*) and homozygous *P. maniculatus* (*m*/*m*); these siblings are hereafter referred to as the “congenic mice”. We tested burrowing behavior in the congenic mice using the sandboxing assay (described above), with three trials per mouse starting at approximately 60 days old. All researchers were blind to the genotypes of the congenic mice.

### Burrowing assay: Tube assay

To identify specific behaviors that contribute to differences in burrow length, we constructed a novel behavioral assay that enabled video recording of mice while they excavated sand (see Fig. 3a), building on our previous assays involving lower resolution video recording of mouse activity in burrows^37^. The new arena consists of a standard housing cage attached to a custom-designed tube extension. The cage and tube are separated by a hand-operated acrylic gate (McMaster-Carr). The tube extension consists of black nylon (3D-printed by Shapeways, New York, NY) with a sheet of clear acrylic on one long side (McMaster-Carr). The 3.8-cm diameter tube has a 4.5 cm long horizontal entryway, followed by 24 cm of a 45° downward sloping tube, which has shallow grooves every 0.5 cm to enable mice to descend and ascend without slipping. We performed all assays under infrared light within 0-4 h of the onset of the mouse’s dark phase and recorded videos with Raspberry Pi Module 2 NoIR cameras (Adafruit) at 30 frames/s.

For each new trial, we prepared the arena by adding a layer of corncob bedding to the housing cage and packing the downward-sloping portion of the tube with wet sand (1:6 volume ratio of sand and water). We placed an individual mouse into the housing cage section of the arena with the gate closed. After 30 min, we manually opened the gate, giving the mouse access to the tube, and video recorded the tube for 60 min. At the conclusion of the trial, we closed the gate and returned the mouse to its home cage. We removed the tube extension and hand-measured the length dug inside the tube to the nearest half centimeter. This assay was repeated three times per mouse, with a minimum of two nights rest between trials. Between each trial, we emptied the tubes and rinsed all arena components with 70% ethanol and water.

### Behavior scoring and analysis

We first identified mouse behaviors through manual observation of *P. maniculatus* and *P. polionotus* tube videos. We identified six behaviors that moved sand (forelimb dig, forelimb push, hindkick, bite/lick, flick, and nose push) and four behaviors that did not (walk, turn around, stationary, and groom; see Table S1 for scoring criteria). To quantify these behaviors, we used BORIS to manually annotate videos and scored these behaviors as point behaviors^38^. For all sand-moving behaviors, a single point represents a single limb/head movement, with the exception of forelimb digging (which consists of rapid forelimb movements anterior to the mouse’s abdomen). Because we could not isolate individual forelimb digging movements at our video frame rate, we scored each instance of forelimb digging with a single point, rather than each individual forelimb stroke.

We also annotated temporal aspects of burrowing behavior. We defined a “visit” to the tube as the duration between when the mouse walks into the tube and when it walks out of the tube (Extended Data Fig. 4a). We defined a digging “bout” as when, during a visit, a mouse executes additional sand-moving behaviors within 1 s of a previous sand-moving behavior (Extended Data Fig. 4b).

For each individual, we scored the videos from only one of the three behavioral trials, the one in which it excavated the most sand. In the case of congenic mice, we scored all videos blind to genotype.

## Acknowledgements

We thank Sade McFadden, Christopher Kirby, Oliver Hollo, Rachelle Ludwick, Sarah Scalia, Julia Mason, and Lynn Shi for their assistance with burrowing assays; Christopher Kirby for assistance creating the congenic mouse line; the Harvard University Center for Brain Science NeuroTechnology Core, Rafael Polo Prieto, and Nina Sokolov for help developing the tube assay. We thank Nicole Bedford, Andres Bendesky, Sandeep Robert Datta, Yuki Haba, Nicholas Jourjine, Lindy McBride, Julius Tabin, Kelsey Tyssowski and Maya Woolfolk for providing helpful feedback on the manuscript. OSH was supported by a National Science Foundation (NSF) Graduate Research Fellowship, a Harvard Quantitative Biology Student Fellowship (DMS 1764269), the Molecular Biophysics Training Grant (NIH NIGMS T32GM008313), an American Society of Mammalogists Grants-in-Aid of Research and a Society for the Study of Evolution R.C. Lewontin Early Award; HCM by a Chapman Memorial Scholarship, a National Science Foundation (NSF) Graduate Research Fellowship, a Doctoral Dissertation Improvement Grant (NSF 1209753), and an American Fellowship from the American Association of University Women; CKH by a Harvard Brain Initiative Young Scientist Transitions Award; EM by a Harvard College Program for Research in Science and Engineering Fellowship; CR by an EU Consortium Joint Masters Scholarship; JIS-S by a Human Frontiers Science Program Postdoctoral Fellowship. HEH is an Investigator of the Howard Hughes Medical Institute.

## Author contributions

OSH, CKH, HCM and HEH conceived of the study. HCM conducted the advanced backcross mapping experiment. OSH performed QTL mapping analyses and created the congenic mouse line. CKH designed the tube assay. CKH and CR measured parental mouse burrowing phenotypes; OSH, CKH, and ELM measured congenic mouse burrowing phenotypes. CKH and JIS-S analyzed behavior data. OSH, CKH, HCM and HEH wrote the manuscript.

## Materials & Correspondence

Correspondence to Hopi E. Hoekstra.

## Data availability

Sequencing data will be uploaded to NCBI SRA. All source data will be included upon publication.

## Code availability

The code used for the analyses will be available from GitHub.

---

**Table S1.** Descriptions of behaviors scored in the tube assay.

| Behavior | Description | Digging behavior? |
| --- | --- | --- |
| Forelimb digging | Alternating/single forelimb strokes that loosen sand. The shoulders will also shift asymmetrically when this occurs. A string of strokes receives a single keystroke from their point of initiation. A string ends when both limbs are placed down in weight-bearing positions or the mouse performs another behavior. | Yes |
| Forelimb Pushing | Both forelimbs simultaneously shift sand underneath the body, towards the hind legs. This is accompanied by arching of the spine with the shoulders in symmetric positions. Every occurrence receives a keystroke. | Yes |
| Hindlimb kick | Hindlimbs simultaneously kick sand behind the mouse. Every occurrence receives a keystroke. | Yes |
| Flick | Single forelimb strokes send sand to the left or right of the body. Every occurrence receives a keystroke. | Yes |
| Nose push | Snout shoves sand forwards. Every occurrence receives a keystroke. | Yes |
| Bite/Lick | Jaws or tongue visibly contact the sand or tube apparatus. Every occurrence receives a keystroke. | Yes |
| Grooming | Limbs and/or mouth are used to clean head or body. A single episode of grooming (which may include grooming of multiple body parts) receives a keystroke. | No |
| Turn around | Mouse rotates such that its anterior and posterior ends swap positions. Rotation that did not result in this swap was not considered. Every occurrence receives a keystroke. | No |
| Stationary | Mouse does not move for one second. Once this time threshold is met, it receives one keystroke. No additional keystrokes are made if the mouse continues to remain still. | No |
| Walking | When entering the tube, the mouse places weight-bearing limbs into the tube. This receives a single keystroke. When exiting the tube, the mouse removes all weight-bearing limbs from the tube. This receives a single keystroke. While in the tube, mouse lifts and sets down each foot in turn, resulting in a movement of the mouse's position. The first step receives a keystroke. | No |

**Table S2.**
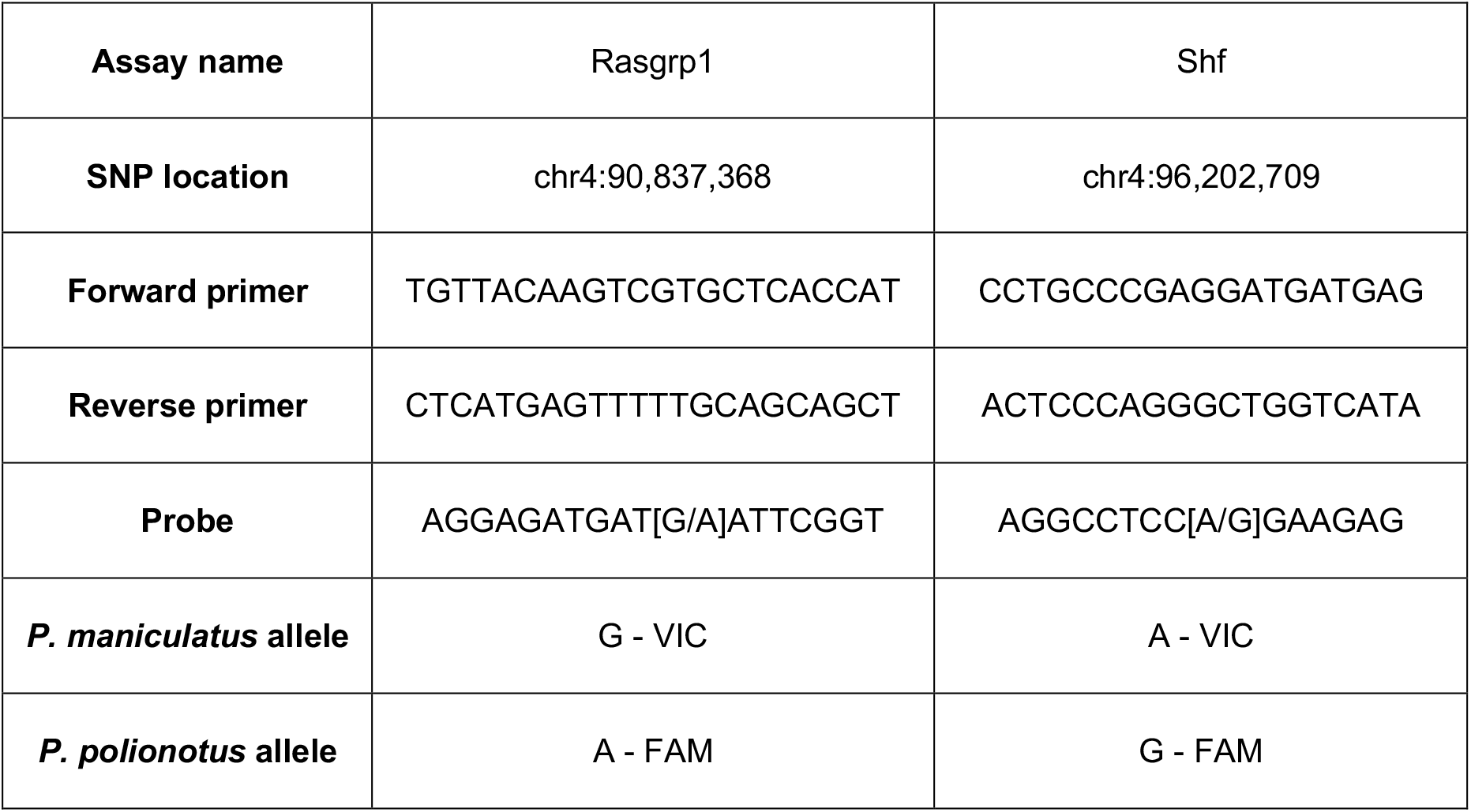
Custom Taqman assay probes used to genotype QTL-chr4 region.

## References

1. Brosset, A. Social organization and nest-building in the forest weaver birds of the genus *Malimbus* (Ploceinae). Ibis 120, 27–37 (1978).

2. Hoikkala, A., Kaneshiro, K. Y. & Hoy, R. R. Courtship songs of the picture-winged *Drosophila planitibia* subgroup species. Anim. Behav. 47, 1363–1374 (1994).

3. Weygoldt, P. Evolution of parental care in dart poison frogs (Amphibia: Anura: Dendrobatidae). J. Zool. Syst. Evol. Res. 25, 51–67 (1987).

4. Tinbergen, N. The hierarchical organization of nervous mechanisms underlying instinctive behaviour. Physiol. Mech. Anim. Behav. 305–312 (1950).

5. Tinbergen, N. The Study of Instinct. (Clarendon Press/Oxford University Press, 1951).

6. Lorenz, K. Z. The evolution of behavior. Sci. Am. 199, 67–82 (1958).

7. Wiltschko, A. B. et al. Mapping sub-second structure in mouse behavior. Neuron 88, 1121– 1135 (2015).

8. Kaplan, H. S., Salazar Thula, O., Khoss, N. & Zimmer, M. Nested neuronal dynamics orchestrate a behavioral hierarchy across timescales. Neuron 105, 562–576.e9 (2020).

9. Seeds, A. M. et al. A suppression hierarchy among competing motor programs drives sequential grooming in Drosophila. eLife 3, e02951 (2014).

10. Hu, C. K. & Hoekstra, H. E. Peromyscus burrowing: a model system for behavioral evolution. Semin. Cell Dev. Biol. 61, 107–114 (2017).

11. Dawson, W. D., Lake, C. E. & Schumpert, S. S. Inheritance of burrow building in Peromyscus. Behav. Genet. 18, 371–382 (1988).

12. Weber, J. N., Peterson, B. K. & Hoekstra, H. E. Discrete genetic modules are responsible for complex burrow evolution in Peromyscus mice. Nature 493, 402–405 (2013).

13. Metz, H. C., Bedford, N. L., Pan, Y. L. & Hoekstra, H. E. Evolution and genetics of precocious burrowing behavior in Peromyscus mice. Curr. Biol. 27, 3837–3845.e3 (2017).

14. Weber, J. N. & Hoekstra, H. E. The evolution of burrowing behaviour in deer mice (genus *Peromyscus*). Anim. Behav. 77, 603–609 (2009).

15. Sumner, F. B. & Karol, J. J. Notes on the burrowing habits of *Peromyscus polionotus*. J. Mammal. 10, 213–215 (1929).

16. Hayne, D. W. Burrowing habits of *Peromyscus polionotus*. J. Mammal. 17, 420–421 (1936).

17. Rand, A. L. & Host, P. Mammal notes from Highland County, Florida. Bull Am Mus Nat Hist 80, 1–21.

18. Xu, M. & Shaw, K. L. The genetics of mating song evolution underlying rapid speciation: linking quantitative variation to candidate genes for behavioral isolation. Genetics 211, 1089–1104 (2019).

19. Ding, Y., Berrocal, A., Morita, T., Longden, K. D. & Stern, D. L. Natural courtship song variation caused by an intronic retroelement in an ion channel gene. Nature 536, 329–332 (2016).

20. Greene, J. S. et al. Balancing selection shapes density-dependent foraging behaviour. Nature 539, 254–258 (2016).

21. Katz, P. S. Evolution of central pattern generators and rhythmic behaviours. Philos. Trans. R. Soc. B Biol. Sci. 371, 20150057 (2016).

22. Savidge, J. A., Seibert, T. F., Kastner, M. & Jayne, B. C. Lasso locomotion expands the climbing repertoire of snakes. Curr. Biol. 31, R7–R8 (2021).

23. Miles, M. C. & Fuxjager, M. J. Synergistic selection regimens drive the evolution of display complexity in birds of paradise. J. Anim. Ecol. 87, 1149–1159 (2018).

24. Wikle, A. W., Broder, E. D., Gallagher, J. H. & Tinghitella, R. M. A rapidly evolving cricket produces percussive vibrations: how, who, when, and why. Behav. Ecol. arad031 (2023).

25. Greenwood, A. K., Wark, A. R., Yoshida, K. & Peichel, C. L. Genetic and neural modularity underlie the evolution of schooling behavior in threespine sticklebacks. Curr. Biol. 23, 1884– 1888 (2013).

26. Jourjine, N. et al. Two pup vocalization types are genetically and functionally separable in deer mice. Curr. Biol. 33, 1237–1248.e4 (2023).

27. Bendesky, A. et al. The genetic basis of parental care evolution in monogamous mice. Nature 544, 434–439 (2017).

28. Layne, J. N. & Ehrhart, L. M. Digging behavior of four species of deer mice (*Peromyscus*). American Museum novitates ; no. 2429. (1970).

29. Bellardita, C. & Kiehn, O. Phenotypic characterization of speed-associated gait changes in mice reveals modular organization of locomotor networks. Curr. Biol. 25, 1426–1436 (2015).

30. Pirger, Z. et al. Interneuronal mechanism for Tinbergen’s hierarchical model of behavioral choice. Curr. Biol. 24, 2018–2024 (2014).

31. Andersson, L. S. et al. Mutations in DMRT3 affect locomotion in horses and spinal circuit function in mice. Nature 488, 642–646 (2012).

32. Peterson, B. K., Weber, J. N., Kay, E. H., Fisher, H. S. & Hoekstra, H. E. Double digest RADseq: an inexpensive method for de novo SNP discovery and genotyping in model and non-model species. PLoS One 7, e37135 (2012).

33. Andolfatto, P. et al. Multiplexed shotgun genotyping for rapid and efficient genetic mapping. Genome Res. 21, 610–617 (2011).

34. Lynch, M. & Walsh, B. Genetics and Analysis of Quantitative Traits. (Sinauer, 1998).

35. Broman, K. W. et al. R/qtl2: Software for mapping quantitative trait loci with high-dimensional data and multiparent populations. Genetics 211, 495–502 (2019).

36. Harringmeyer, O. S. & Hoekstra, H. E. Chromosomal inversion polymorphisms shape the genomic landscape of deer mice. *Nat*. Ecol. Evol. 6, 1965–1979 (2022).

37. Bedford, N. L. et al. Interspecific variation in cooperative burrowing behavior by Peromyscus mice. Evol. Lett. 6, 330–340 (2022).

38. Friard, O. & Gamba, M. BORIS: a free, versatile open-source event-logging software for video/audio coding and live observations. Methods Ecol. Evol. 7, 1325–1330 (2016).

